# Principles Underlying the Input-Dependent Formation and Organization of Memories

**DOI:** 10.1101/346478

**Authors:** Juliane Herpich, Christian Tetzlaff

**Author notes:** Member of the International Max Planck Research School for Physics of Biological and Complex Systems at the University of Göttingen, IMPRS-PBCS, Germany.

## Abstract

The brain of higher-order animals continuously and dynamically learns and adapts according to variable, complex environmental conditions. For this, the neuronal system of an agent exhibits the remarkable ability to dynamically store and organize the information of plenty of different environmental stimuli into a web of memories, which is essential for the generation of complex behaviors. The basic structures of this web are the functional organizations between two memories such as discrimination, association, and sequences. However, how these basic structures are formed robustly in an input-dependent manner by the underlying, complex and high-dimensional neuronal and synaptic dynamics is still unknown. Here, we develop a mathematical framework which reduces the complexity in several stages such that we obtain a low-dimensional mathematical description. This provides a direct link between the involved synaptic mechanisms, determining the neuronal and synaptic dynamics of the network, with the ability of the network to form diverse functional organizations. Given two widely-known synaptic mechanisms, correlation-based synaptic plasticity and homeostatic synaptic scaling, we use this new framework to identify that the interplay of these mechanisms enables the reliable formation of sequences and associations between two memory representations - missing the important functional organization of discrimination. We can show that this shortcoming can be compensated by considering a third mechanism as inhibitory synaptic plasticity. Thus, the here-presented framework and results provide a new link between diverse synaptic mechanisms and emerging functional organizations of memories. Furthermore, given our mathematical framework, one can now investigate the principles underlying the formation of the web of memories.

**AUTHOR SUMMARY:** Higher-order animals are permanently exposed to a variety of environmental inputs which have to be processed and stored such that the animal can react appropriate. Thereby, the ongoing challenge for the neuronal system is to continuously store novel and meaningful stimuli and, dependent on their content, to integrate them into the existing web of knowledge or memories. The smallest organizational entity of such a web of memories is described by the functional relation of two interconnected memories: They can be either unrelated (discrimination), mutually related (association), or uni-directionally related (sequence). However, the neuronal and synaptic dynamics underlying the formation of such structures is mainly unknown. To investigate possible links between physiological mechanisms and the organization of memories, in this work, we develop a general mathematical framework enabling analytical approach. Thereby, we show that the well-known mechanisms of synaptic plasticity and homeostatic scaling in conjunction with inhibitory synaptic plasticity enables the reliable formation of all basic relations between two memories. This work provides a further step in the understanding of the complex dynamics underlying the organization of knowledge in neural systems.

## INTRODUCTION

Learning and memorizing various pieces of information from the environment are vital functions for the survival of living beings. In addition, the corresponding neuronal system has to learn the environmental relations between these different pieces. For this, the neuronal system has to form memory representations of the information and to organize them accordingly. However, the neuronal and synaptic dynamics determining the organization of these representations are widely unknown.

The synaptic-plasticity-and-memory hypothesis relates the formation of memory representations to the underlying neuronal and synaptic mechanisms [1, 2]. Namely, a to-be-learned piece of information activates via an environmental stimulus a certain population of neurons triggering synaptic plasticity. Synaptic plasticity, in turn, changes the weights of the synapses between the activated neurons such that these neurons become strongly interconnected and form a memory representation – so-called Hebbian cell assembly (CA) – of the presented information [3–6]. Besides the formation of a memory representation, the newly learned piece of information is also related to already stored information [3, 7–9]. Thereby, the relations or functional organizations between different memory representations can be organized in three different, fundamental ways: they can be unrelated (discrimination), mutually related (association), or uni-directionally related (sequence). However, although the link between the formation of a single memory representation and the underlying neuronal and synaptic mechanisms is already well established [10–13], it is largely unknown which mechanisms enable the self-organized formation of relations *between* memory representations.

In this theoretical study, we have developed the first theoretical framework enabling to analyze the ability of diverse neuronal and synaptic mechanisms to form memory representations and, in addition, to form the different types of memory-relations. Thereby, our analysis indicates that the interaction of correlation-based synaptic plasticity with homeostatic synaptic scaling is not sufficient to form all types of memory-relations, although it enables the formation of individual memory representations [11, 14]. However, our analysis shows that, if the average level of inhibition within the memory representations is significantly lower than the average level in the remaining network, the neuronal system is able, on the one hand, to form memory representations and, on the other hand, to organize them into the fundamental types of memory relations in an input-dependent, self-organized manner.

Several theoretical studies [11–15] investigated the formation of individual memory representations in neuronal systems indicating correlation-based synaptic plasticity as essential mechanism. In addition, homeostatic plasticity, as synaptic scaling [16], is required to keep the system in an adequate dynamic regime [17–19]. Further studies indicate that synaptic plasticity and homeostatic plasticity also yield the formation of sequences of representations [15, 20, 21]. However, it remains unclear whether the interaction of synaptic and homeostatic plasticity also enables the formation of further memory-relations described before. Interestingly, several theoretical studies [7, 22, 23] indicate that a neural system with the ability to form all described memory-relations has an algorithmic advantage to process the stored information. Furthermore, the neuronal dynamics resulting from interconnected memory representations match experimental results on psychological [24] and single-neuron level [25, 26]. However, these studies consider neural systems after completed learning; thus, it is unclear how neuronal systems form the required relations between memory representations in a self-organized manner.

We consider a neuronal network model with plastic excitatory connections, which are governed by the interaction of correlation-based and homeostatic plasticity. As already shown in previous studies, this interaction enables the self-organized formation of individual memory representations [11, 14]. We analyze the ability of the plastic network to form different types of relations between two memory representations – namely, discrimination, sequences, and association. Please note that this is a high-dimensional problem of the order of *N*^2^ (given *N* neurons). To reduce complexity, standard approaches as mean-field analysis are not feasible, as they obliterate the different memory representations involved. Thus, we developed a new theoretical framework by considering the mean equilibrium states of the relevant system variables and by comparing them to constraints given for the different memory-relations. Thereby, we map the constraints on the long-term average activity level of the neuronal populations involved reducing the problem to a 2-dimensional one, which can be analyzed graphically and analytically. By this framework, we optimized the parameters of the system and identified that correlation-based and homeostatic plasticity do not suffice to form all three types of memory-relation. Instead, if the average inhibitory level within the memory representations is below control level, memory representations can be formed, maintained and related to each other. In addition, we show that the required state can also be reached in a self-organized, dynamic way by the interplay between excitatory and inhibitory synaptic plasticity. Thus, the here-presented results provide a next step to understand the complex dynamics underlying the formation of memory relations in neuronal networks.

## RESULTS

In our work, we analyze the ability of two neuronal populations *p* ∈ {1, 2} to become memory representations and, in parallel, to reliably build up different functional organizations such as discrimination, sequence, and association (Tab. I).

**Table I.** Synaptic weight-dependent definition of memory and different forms of functional organization of two interconnected neuronal populations.

| memory representation | $\langle \tilde{\omega}_{11} \rangle$ | $\langle \tilde{\omega}_{22} \rangle$ | $\langle \tilde{\omega}_{21} \rangle$ | $\langle \tilde{\omega}_{12} \rangle$ | functional organization | abbreviation | color code |
| --- | --- | --- | --- | --- | --- | --- | --- |
| **x** | $< \tilde{\theta}$ | $< \tilde{\theta}$ | - | - | none | nm | grey |
| **✓** | $> \tilde{\theta}$ | $> \tilde{\theta}$ | $< \tilde{\theta}$ | $< \tilde{\theta}$ | discrimination | disc | blue |
| | | | $< \tilde{\theta}$ | $> \tilde{\theta}$ | sequence 12 | s12 | yellow |
| | | | $> \tilde{\theta}$ | $< \tilde{\theta}$ | sequence 21 | s21 | green |
| | | | $> \tilde{\theta}$ | $> \tilde{\theta}$ | association | asc | red |
|  |  |  | bistable |  | various | bs | pink |

The different functional organizations are defined based on the average excitatory synaptic weights $\langle \tilde{\omega}_{11} \rangle$ , $\langle \tilde{\omega}_{22} \rangle$ , $\langle \tilde{\omega}_{21} \rangle$ , and $\langle \tilde{\omega}_{12} \rangle$ at equilibrium in relation to the average inhibitory synaptic weights ( $\tilde{\theta}$ ).

**Table II.**
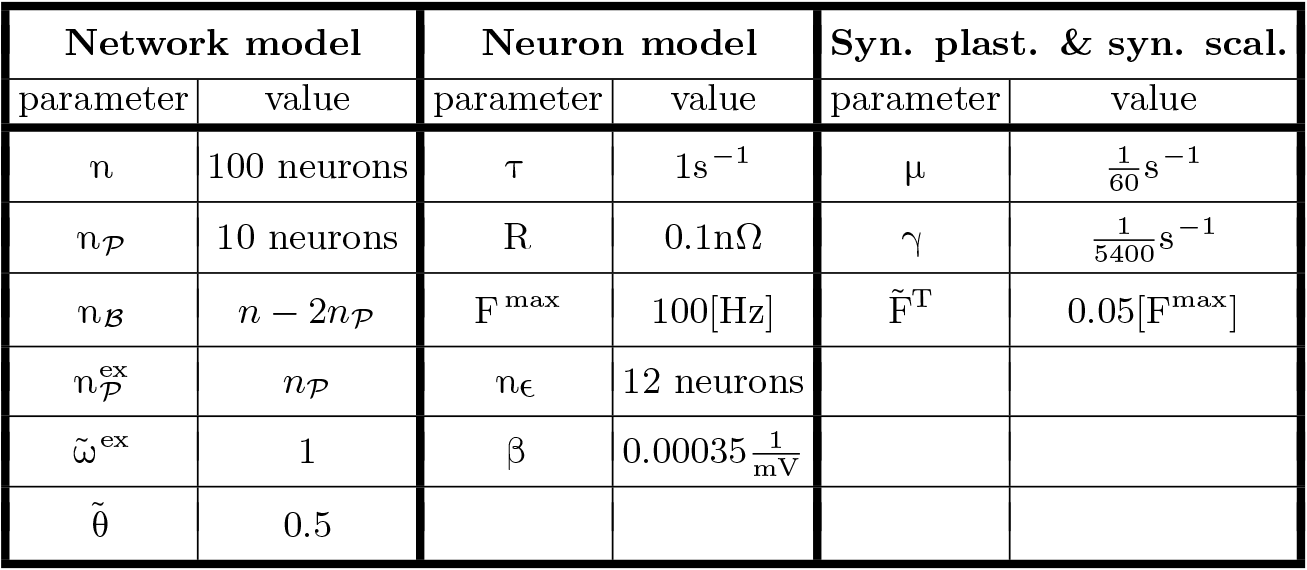
Used Parameters.

In general, the external input to population *p* should trigger synaptic changes within the population such that it becomes a memory representation of its specific input. Given two of these populations, dependent on the input properties, connections between the populations should also be altered to form the neuronal substrate underlying the diverse functional organizations described before. In accordance to the synaptic-plasticity-and-memory hypothesis [1, 3], we define a neuronal population as being a *memory representation* if its neurons are strongly interconnected. Thus, the average excitatory synaptic strength between all neurons within the population has to be larger than the average inhibitory synaptic strength. Thus, due to the dominant excitation, neuronal activity within the population will be amplified. We define the relation between two memory representations in a similar manner based on the relation of excitation and inhibition between the corresponding neuronal populations: In general, if the average excitatory synaptic strength from one population to the other is larger than the average inhibitory synaptic strength, an increased level of activity in the former population triggers an increased activation in the latter. This can be different for both directions such that, for instance, the net connection from population 1 to 2 can be excitatory and inhibitory from 2 to 1. This case is defined as a *sequence* from 1 to 2. Similar, an *association* is present if both connections are excitatory-dominated, and a *discrimination* consists of both directions being zero or inhibition-dominated.

To analyze the self-organized formation of memory representations and their functional organization, we consider a plastic recurrent neuronal network model 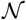 consisting of rated-coded neurons being interconnected via plastic excitatory and static inhibitory connections (Fig 1 A). Within the recurrent network are two distinct populations of neurons (*p* ∈ {1, 2}; black and yellow dots, respectively) within each the neurons receive the same external input 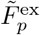(red layer *ε*). All remaining neurons are summarized as background neurons 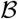 (blue) such that the neuronal network can be described as the interaction of three different neuronal populations.

**Fig 1.**
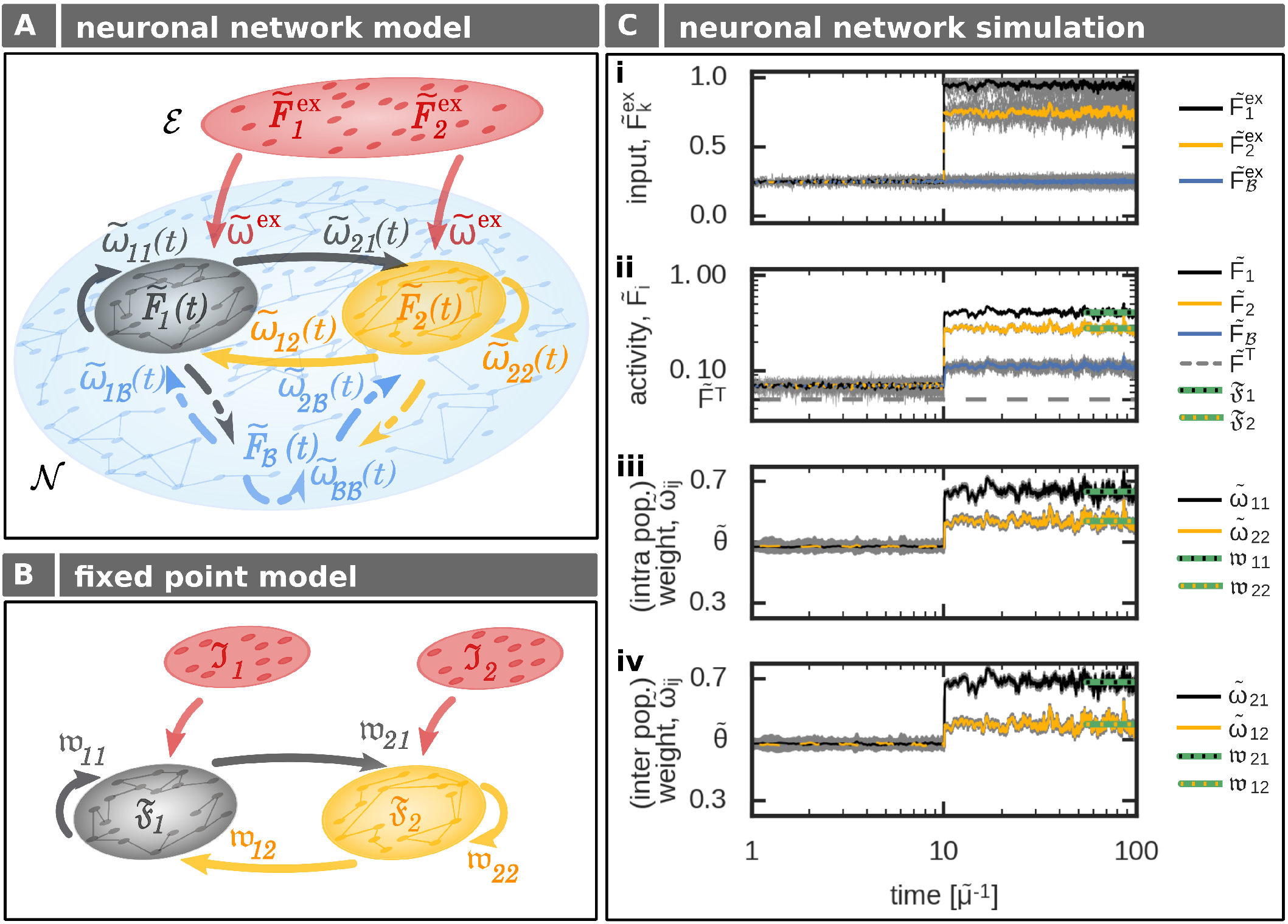
The formation of interconnected memory representations in a plastic neural network. (**A**) In a recurrent network 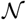 two neuronal populations (1: black; 2: yellow) receive specific external inputs of average amplitudes 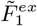 and 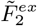. All remaining neurons of the network (blue) are combined to a background population 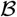 and serve as control neurons receiving noisy external inputs. Each population 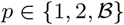 is described by its mean intra-population-synaptic weight 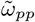, its mean activity 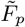, and its connections to other populations 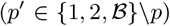 via a set of synapses with average synaptic strength 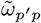. (**B**) The abstraction of the neuronal network model yields a low-dimensional one described by the mean equilibrium activities (𝔉_*p*_) and corresponding mean equilibrium synaptic weights (𝔴_*p*′*p*_). Here, the external input (red) combines inputs from background neurons and external inputs given in the complete network model (A). (**C**) In the network model (A), changing the amplitude of the external input to neuronal populations 1 (black) and 2 (yellow) at *t* = 10 yields increased average activities within the populations and background neurons (blue) triggering synaptic changes. Gray lines indicate single neuron/synapse dynamics. After a brief period, all system variables reach an equilibrium state. This state is matched very well by the theoretical analysis (green lines) considering the abstract model (B). (i) inputs; (ii) average activities of each population; (iii) average intra-population synaptic weights; (iv) average inter-population synaptic weights. The average input amplitudes are determined by two Ornstein-Uhlenbeck processes with mean 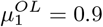 and 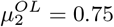.

All excitatory connections within the recurrent layer are plastic regarding the interaction of fast correlation-based synaptic plasticity and slow homeostatic synaptic scaling [18, 27]. Please note that previous studies indicate that this interaction yield the reliable formation of individual memory representations [11, 14]. The resulting changes in synaptic weights between postsynaptic neuron *i* and presynaptic neuron *j* is thus regulated by

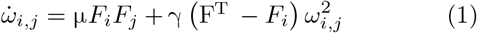

with neural activities *F*_*i*_ and *F*_*j*_, μ being the time scale of synaptic plasticity, γ the time scale of synaptic scaling, and the target firing rate F^T^ of the homeostatic process.

Thus, an external input to populations 1 and 2 alters neural activities within the corresponding populations and, furthermore, triggers changes in the corresponding synaptic weights (see Fig 1 C for an example). In the first phase, all neurons of the network receive a noisy input (Fig 1 C, Panel i) such that neural activities (Panel ii) and synaptic weights (Panels iii and iv) are at base level. At *t* = 10, both populations 1 and 2 receive a strong external input (Panel i). In more detail, each neuron in a specific population receives an input from 10 input neurons each modeled by its own Ornstein-Uhlenbeck process (grey lines; yellow and black line indicate the average). The mean of these processes is the same for all input neurons transmitting to one population (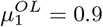 for pop. 1 and 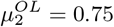 for pop. 2). After a brief transition phase, the system reaches a new equilibrium state. Here, for both populations the intra-population synapses are stronger than the average inhibitory synaptic weights (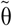; Panel iii) indicating the formation of two memory representations. Furthermore, the excitatory synapses connecting both populations are adapted and also become stronger than the average inhibition level (Panel iv). This implies that both populations or memory representations are strongly linked with each other; thus, an association has been formed. Therefore, given a certain stimulus, the equilibrium state of the synaptic weights determines the functional organization of the corresponding memory representations.

### Memory representation and functional organization

As the impact of single synapses on the overall network dynamics is small, we will consider in the following the equilibrium states of the average synaptic weights of inter- and intra-population synapses (indicated by 〈*x*〉). Thus, these synaptic states determine whether a neuronal population is a memory representation, and how several of these representations are functionally organized (discrimination, sequence, or association).

Therefore, as long as the average recurrent, or intrapopulation excitatory synaptic weight 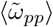 of neuronal population *p* is weaker than the average inhibitory synaptic strength 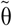, an external input to the population will lead to an average decrease in population-activity (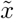 indicates the normalized variable of *x*; see Methods). Thus, the neuronal population does not serve as a memory representation and the system state is defined as *no memory state* (*nm*; Tab. I, Fig 2 Ai, left panel, grey area). By contrast, if the average recurrent excitatory synaptic weight is at the equilibrium state above the level of inhibition (Fig 2 Ai, left panel, white area), the neuronal population reacts with an increased activity level to an external input and, therefore, it serves as a memory representation (*memory state*). In other words, the neuronal population *p* has to fulfill the following condition to be classified as a memory representation:

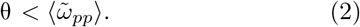

**Fig 2.**
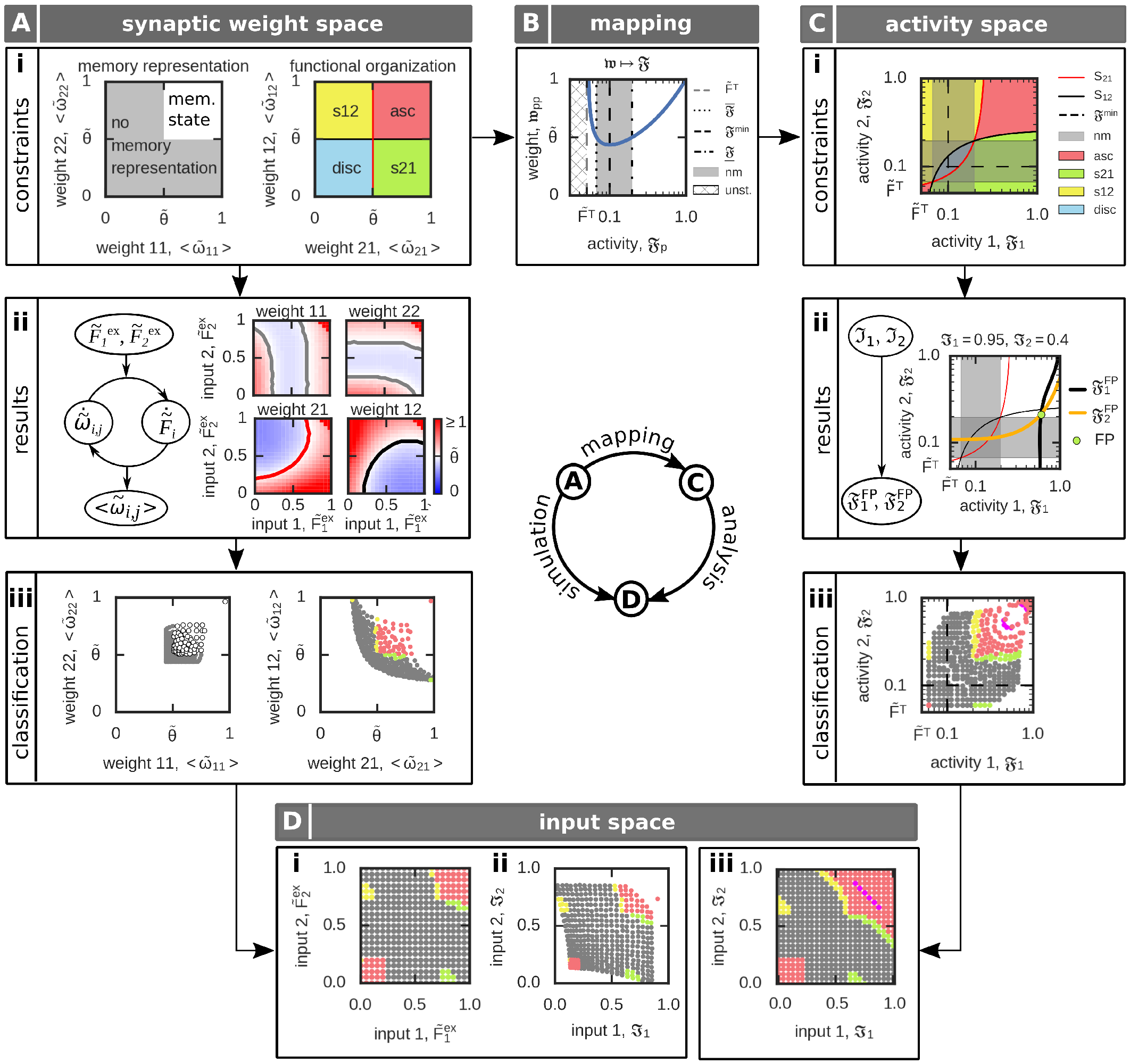
Definition and analysis of the input-dependent formation of functional organizations (FO) between two neuronal populations. (**A i**) The different FOs are defined based on the average synaptic weights. Details see main text. (**A ii**) Solving numerically the complete *N*^2^-dimensional network dynamics for different external inputs (left) yields average excitatory synaptic weights. (**A iii**) These average synaptic weights (A ii) can be analyzed regarding the weight-dependent conditions of FOs (A i). (**B**) Considering the dependency of the intra-population synaptic weight (𝔴_*pp*_, blue curve) on its respective activity (𝔉_*p*_) in equilibrium enables the mapping of the weight-dependent memory conditions on the 2*d*-activity space. grey space: no memory representation; white space: memory representation. (**C i**) Conditions for the different FOs of two memories in the mapped 𝔉_1_ − 𝔉_2_–activity space of the neuronal populations. (**C ii**) Within this 2*d*-space one can calculate the fixed point of the population activities 𝔉_1_, 𝔉_2_ by the intersection of the equations 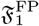 and 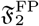 given an input stimulation ℑ_1_, ℑ_2_. (**C iii**) By comparing the resulting fixed point from (C ii) with the FO-conditions (C i), we can obtain the respective FO. (**D**) Given the results in the weight- (A) or activity-space (B), we can assess for each input case the resulting FO. Used parameters: 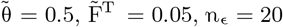.

Given that both neuronal populations *p* ∈ {1, 2} are a stable memory representation (Fig 2 Ai, left panel, white area), they can form different functional organizations (discrimination, sequences, or association). Thereby, the average inter-population synaptic weights 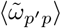 (*p*,*p*′ ∈ {1, 2}, *p* ≠ *p*′) define the different functional organizations, dependent on their relation to the average inhibitory synaptic weight strength 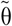 (Tab. I). Thus, for two interconnected memories 1 and 2, we can define four different functional organizations with different weight-dependent conditions (Fig 2 Ai, right panel):

- *discrimination*: both average inter-population synaptic weights are weaker than the inhibitory weights (blue, *disc*)

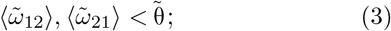
- *sequence 21*: average inter-population synaptic weight from memory 1 to memory 2 is stronger than inhibitory weights, while the inter-population synaptic weight from 2 to 1 is weaker (green, *s21*)

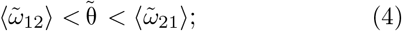
- *sequence 12*: inter-population synaptic weight from memory 1 to memory 2 is weaker than the inhibitory weights, while the inter-population synaptic weight from 1 to 2 is stronger (yellow, *s12*)

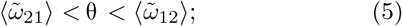
- *association*: both average inter-population synaptic weights are stronger than the average inhibitory synaptic weight (red, *asc*)

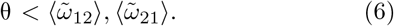

#### Full-network analysis

Assessing under which input-condition the plastic neuronal network is able to form memory representations and diverse functional organizations, the whole set of *N*^2^ + 2 · *N* differential equations has to be solved numerically for each input condition 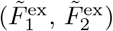. Thereby, each simulation runs until the system reaches its equilibrium state. In this equilibrium state, excitatory synaptic weights are analyzed and compared to the inhibitory synaptic weights (Fig 2 A ii, right panels) enabling a classification according to the functional organizations (Fig 2 A iii). This classification can be mapped to the inputs providing the resulting functional organization dependent on the specific external inputs (Fig 2 D i,ii). Note, for better comparison with the population model (see the next section), the results (Fig2 D i) are mapped to the population-input-space defined below (Fig 2D ii). The whole analysis is computational expensive and, furthermore, it does not provide additional insights into the relation between the synaptic dynamics and the ability to form diverse functional organizations. Thus, in the following, we provide a different approach to solve this complex, high-dimensional mathematical problem.

#### Population model at equilibrium

To reduce the complexity of the system, in the following, we derive a method that directly calculates the mean state variables of the memory-related neuronal populations *p* ∈ {1, 2} at equilibrium (Fig 1 B). For this we combine the inputs a population receives from the external layer *ε* 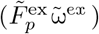 with the inputs from the background neurons in 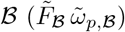 to

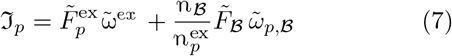

with 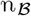 being the number of neurons belonging to the background population 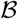 and 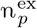 being the number of input neurons.

Given this input stimulation ℑ_*p*_, we consider that the firing rate 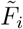 of each neuron *i* ∈ *p* of a population *p* is close to the mean firing rate of this particular population 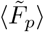. By this, we receive the mean neuronal activity of population *p* at equilibrium 𝔉_*p*_ (Eq 29). With the equilibrium activities of the neuronal populations *p* and *p*′, in turn, we can calculate the respective equilibrium synaptic weights 𝔴_*p*′*p*_ from population *p* to population *p*′ (Eq 30):

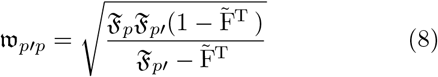

and the equilibrium synaptic weights of each population *p* itself

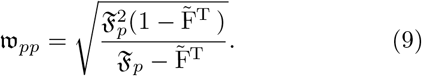

#### Activity-dependent constraints of memory representation and functional organization

Given the relation between average population activities and synaptic weights in equilibrium (Eqs 8 and 9), next, we map the weight-dependent conditions for memory representations (Eq 2) and functional organizations (Eqs 3-6) onto the average population activities (Fig 2 B). Thus, the fixed point or equilibrium equation of the synaptic dynamics (Eq 9; blue curve in Fig 2 B) yields two activity-dependent conditions of a neuronal population *p* to become a memory representation:

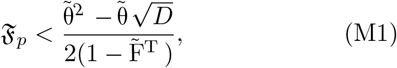

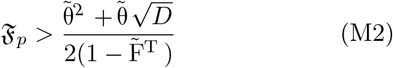

with 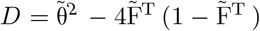. Thus, we can define two open intervals (Fig 2 B, white regimes) for the population activity leading to a representation of the respective memory by:

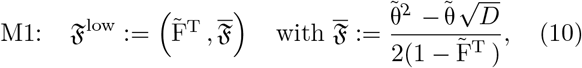

and

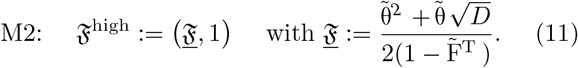

Note that below the target firing rate 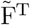 the interaction of synaptic plasticity and scaling does not have a fixed point (Fig 2 B, hatched regime; Eq 1). Furthermore, in the regime 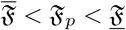 no proper memory representation can be formed (equivalent to 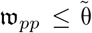). This activity regime is defined as the no memory state nm:= 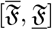 (Fig 2 B, grey regime) with size 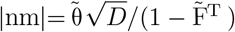.

As we consider the interaction of two interconnected neuronal populations 1 and 2, we receive four distinct activity regimes enabling the formation of two memory representations (Fig 2 C i). These regimes are defined by all possible combinations of 𝔉^low^ and 𝔉^high^ in both dimension of 𝔉_1_ and 𝔉_2_. In other words, these activity regimes are separated by the no memory phase (nm) in both dimensions (Fig 2 C i, grey regimes).

Similar, with Eq 8, we can map the diverse conditions of the functional organizations (Eqs 3-6) onto different activity-dependent conditions. In general, the condition 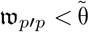 becomes

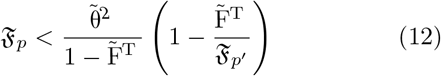

and 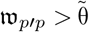 to

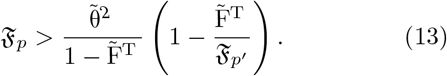

To distinguish between both cases (which determines the functional organization between two memories 1 and 2), we define two separatrixes:

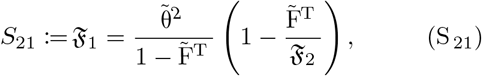

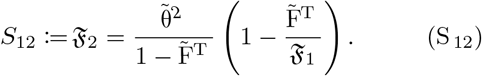

Thus, *S*_21_ represents 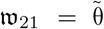 in the activity-space (Fig 2 A i, C i, red curves) while S_12_ represents 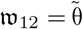 (Fig 2 A i, C i, black curves).

##### Discrimination

When both activities 𝔉_1_ and 𝔉_2_ are below the respective separatrix *S*_21_ and *S*_12_ (Fig 2 C i, blue regime), the system is in an discriminatory functional organization.

##### Sequence

The system establishes a sequence from memory 1 to memory 2, when the activity 𝔉_1_ is above the corresponding separatrix *S*_21_ while the activity 𝔉_2_ stays below separatrix *S*_12_ (Fig 2 C i, green, s21) and vice versa for a sequence from memory 2 to memory 1 (yellow, s12).

##### Association

Both memories are organized in an associational entity when both neuronal activities 𝔉_1_ and 𝔉_2_ are above their respective separatrix (Fig 2 C i, red phase).

#### Functional organizations in activity-space

To obtain which functional organization the system forms for a given external input, we have to calculate the input-dependent average population activities 𝔉_1_ and 𝔉_2_ in the equilibrium state. For this, for each pair of inputs ℑ_1_ and ℑ_2_, we derive the fixed point conditions for both populations (Eq 34) dependent on the activity of the other population (Fig 2 C ii; 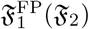, black curve; 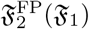, yellow curve). The intersection between both fixed point conditions 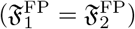 determines the fixed point of the whole system (green dot). The relation of the corresponding activities 𝔉_1_ and 𝔉_2_ of this intersection to the separatrixes determines the functional organization (Fig 2 C iii). This can be expressed in the input space (Fig 2 D iii). Thus, the interaction of synaptic plasticity and scaling enables the formation of sequences in both directions and associations. Furthermore, there is a regime of input values in which no memory representation is formed.

Comparing the analytical results from the population model (Fig 2 D iii) with the results from the full network analysis (Fig 2 D ii) indicates that the population model matches the full network quite well. Especially, the inherent property of a system to form different functional organizations is precisely predicted by the population model. Remarkably, already the mapping of the weight-dependent conditions on the activity-space (Fig 2 C i) provides sufficient information to assess the possible organizations of memories for a given system (not requiring the evaluation of the system’s fixed points).

#### Synaptic-plasticity-induced formation of associations

Both analysis methods (Fig 2 A and C) indicate that the interaction of synaptic plasticity and scaling enables the formation of sequences and associations between two memory representations (Fig 2 D). Interestingly, for very low external input stimulations ℑ_1_ and ℑ_2_, the system forms an association. This is mainly due to the quadratic weight-dependency of synaptic scaling (Eq 1) such that for low population activities synaptic scaling dominates the synaptic dynamics and drives the synaptic weights to high values (up-scaling). This is in contrast to the synaptic-plasticity-and-memory hypothesis [1, 3], which states that the processes of correlation-based synaptic plasticity dominates learning. Along this line, synaptic scaling should mainly regulate the synaptic dynamics in an homeostatic manner [28]. In other words, the synaptic weights should mainly increase with increasing neuronal activities. To determine the regime in which this behavior is present, we consider the activity level 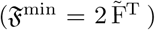 yielding the local minimum of the synaptic weight function (Eq 9; Fig 2 B). Below 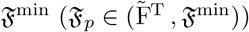, the dynamics are dominated by synaptic scaling and should be avoided. Thus, the plausible activity regime is in the

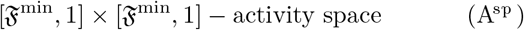

(Fig 3 Ai, blue space) as in this space the synaptic weight dynamics are dominated by correlation-based synaptic plasticity. This regime exists as long as the target activity 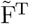 is below 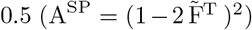 restricting the target activity parameter to 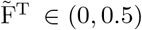.

**Fig 3.**
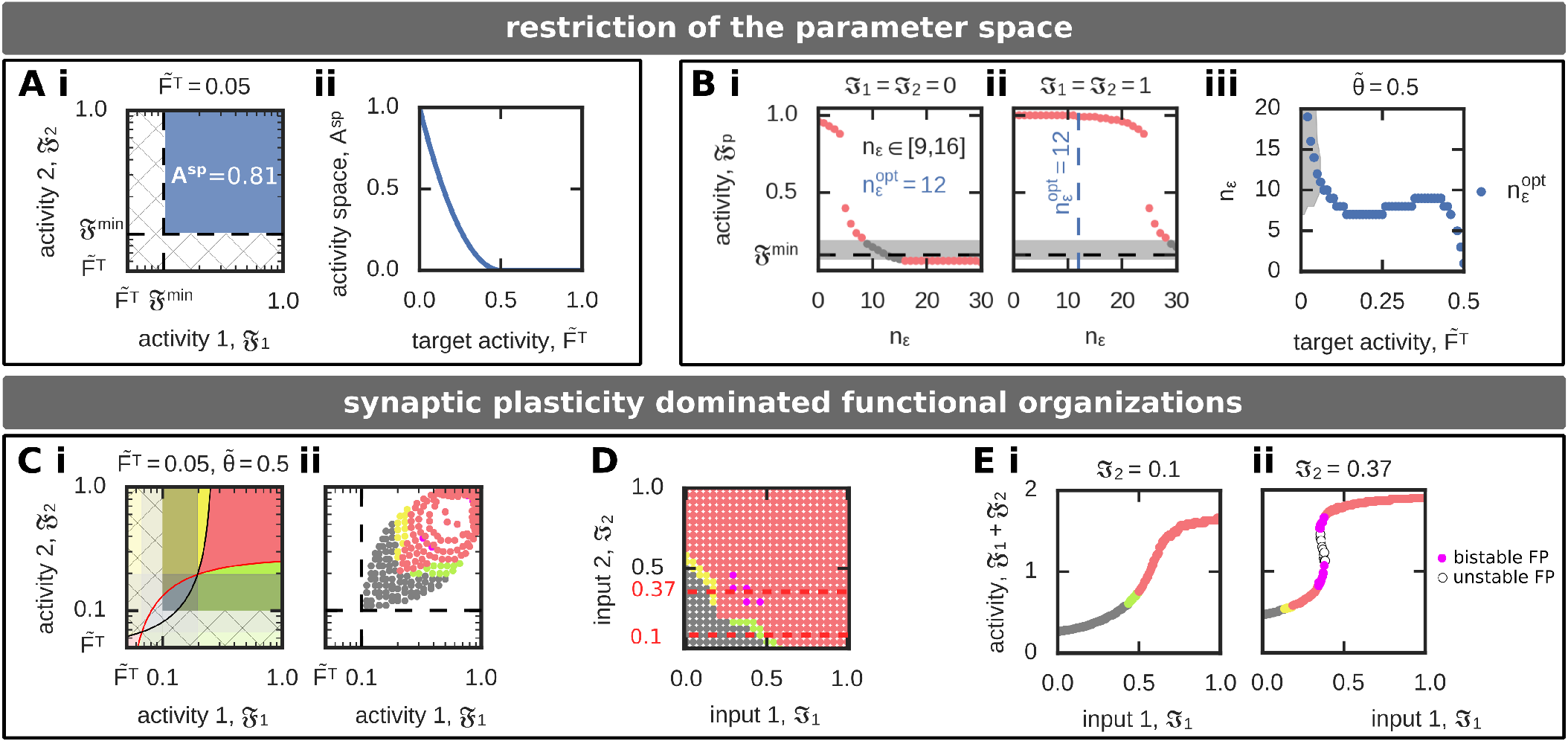
Synaptic plasticity dominated functional organization (FO) of two interconnected memories. **(A, B)** The regime A^SP^, in which synaptic plasticity dominates the synaptic dynamics, depends on the target firing rate 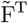 (A) and inflexion point 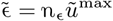 (B). **(A)** The area of the 𝔉_1_ − 𝔉_2_–activity phase space (A i, blue space) leading to synaptic plasticity dominated FOs decreases with increasing target firing rate 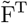 (A ii). (**B**) The inflexion point (measured in *n*_ϵ_) determines the activity-input mapping such that for the same input different activities and, thus, different FOs are realized. If the inflexion point equals *n*_ϵ_ ^opt^, 𝔉_*p*_ ≈ 𝔉^min^. (**B i**): ℑ_1_ = ℑ_2_ = 0. (**B ii**): ℑ_1_ = ℑ_2_ = 1. (**B iii**) The value of *n*_*ϵ*_ ^opt^ (blue) depends on the target firing rate 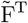. The grey area specifies all *n*_*ϵ*_ that yield to the no-memory state. (**C - E**) One example of synaptic plasticity dominated formation of FOs. Used parameters: 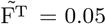, *n*_*ϵ*_ = 12. (*C*) Although the system implies regimes of scaling-dominated synaptic dynamics (hatched area; i), the activity-input mapping excludes that the system can reach these by external inputs (ii). (**D**) The resulting ℑ_1_ − ℑ_2_–input phase space, color-coded according to the respective FOs, shows that associations can only be formed for stronger inputs (compare to Fig 2 D iii). (**E**) The sum of both population activities (𝔉_1_ + 𝔉_2_) for a fixed input ℑ_2_ shows for some cases the existence of two equilibrium states encoding associations (pink). (**E i**): ℑ_2_ = 0:1. (**E ii**): ℑ_2_ = 0:37.

##### Theorem 1

The activity regime 𝔉^low^ ((M1), Eq 10), which enables a proper formation of memory representations, is not part of the correlation-based dominated activity space (A^sp^) for synaptic plasticity.

##### Proof.

Assume that the upper bound 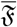 of 𝔉^low^ is smaller than the lower bound 𝔉^min^ for the synaptic plasticity dominated activity regime (A^*sp*^). It follows that the condition 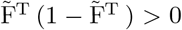 is true for 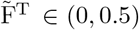 and by this 𝔉^low^ ∉ A^sp^.

Thus, to assure that the activity regime 𝔉^low^ cannot be reached by the system, we have to change the mapping between neuronal activity and inputs such that no reasonable input pair ℑ_1_ ℑ_2_ yields population activities within 𝔉^low^. This activity-input mapping is mainly determined by the inflexion point ϵ of the activity function (Eqs 24 and 34). Here, we specify the inflexion point in units of *n*_ϵ_ (ϵ = *n*_ϵ_*u*^max^) with *n*_ϵ_ being the number of maximally active pre-synaptic neurons (with maximally strong synapses, see Methods). First, we analyze the resulting population activities 𝔉_*p*_ and corresponding functional organizations for different *n*_ϵ_ given no external inputs (ℑ_1_ = ℑ_2_ = 0; Fig 3 B i). For *n*_ϵ_ > 12, the population activities are below 𝔉^min^, which triggers up-scaling yielding a scaling-induced formation of an association. Please note that the system analyzed beforehand (Fig 2) has *n*_ϵ_ = 20. For *n*_ϵ_ < 9, neurons are too easy to excite such that activities are independent of the input nearby the maximum yielding the functional organization of association. For 9 ≤ *n*_ϵ_ ≤ 16, the system is in the no memory state. Thus, to prevent the input-*in*dependent association of two interconnected neuronal populations, we consider the inflexion point to be in the regime 9 ≤ *n*_ϵ_ ≤ 16. Thereby, *n*_ϵ_^opt^ = 12 yields activities nearby the lower minimum activity level 𝔉^min^ defined above. The same analysis for maximal external input stimulation (ℑ_1_ = ℑ_2_ = 1; Fig 3 B ii) shows that for *n*_ϵ_^opt^ = 12 the system can nearly reach its maximal firing rate of 𝔉_*p*_ = 1 such that the whole activity space 𝔉^min^ − 1 can be reached by the system. Please note that for *n*_ϵ_ > 24 the system cannot reach high activity levels and for *n*_ϵ_ > 27 the system is not able to form memory representations, although it is maximally stimulated by the external input. Furthermore, as can be expected from Fig 3 A, the value of *n*_ϵ_^opt^ depends on the target firing rate 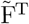 (Fig 3 B iii).

In the following (Fig 3 C-E), we will consider *n*_ϵ_ = 12 and 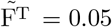, which implies that an association is only be formed by synaptic dynamics dominated by correlation-based synaptic plasticity. The activity regime yielding scaling-dominated learning (hatched area in Fig 3 C i) is theoretically possible, however, the adapted activity-input mapping assures that this regime cannot be reached for given external inputs (Fig 3 C ii and D). In the resulting system, low inputs ℑ_1_,ℑ_2_ lead to a no-memory state (grey), while in a small regime sequences are formed (yellow and green). Thereby, the sequence is formed from the population receiving a stronger input to the population receiving the weaker input. If both inputs are strong, an association between the memory representations is being build (red). Note that there is a small bimodal regime with two long-term equilibrium states both being an association (pink; see two exemplary cross sections in Fig 3 E).

#### Parameter-dependency of functional organizations

After optimizing the activity-input mapping by *n*_ϵ_ such that the formation of diverse functional organizations is dominated by synaptic plasticity, in the following, we will analyze which kind of functional organizations can be formed by the system dependent on the different system parameters. Thereby, we will focus on the target activity 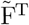 and the average level of inhibition 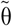. In general, as the no-memory state implies that neuronal populations can exist which do not encode information (or have “forgotten” this information), this state has a large influence on the overall system properties. As described before, the size of this state is given by 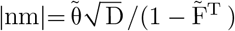 with 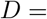 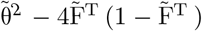. Thus, the discriminant *D* or rather 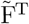 and 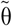 define whether the no-memory state can exist in a given system (Fig 4). In addition, as the relation of the activity levels to the separatrices S_21_ and S_12_ define which kind of functional organization is present (see above), the separatrices have to be within the synaptic plasticity dominated activity-regime (𝔉_*p*_ ∈ (𝔉^min^, 1)) to enable the formation of sequences (*s12*: 𝔉_1_ < S_21_, 𝔉_2_ > *S*_12_; *s21*: 𝔉_1_ > *S*_21_, 𝔉_2_ < *S*_12_) and discrimination (*disc*: 𝔉_1_ < *S*_21_, 𝔉_2_ < *S*_12_). As *S*_12_ (*S*_21_) increase with 𝔉_1_ (𝔉_2_), the maximum difference *S* between the lower activity level of synaptic plasticity dominated dynamics and the separatrix is given for 𝔉_1_ = 1 (𝔉_2_ = 1) such that

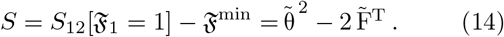

**Fig 4.**
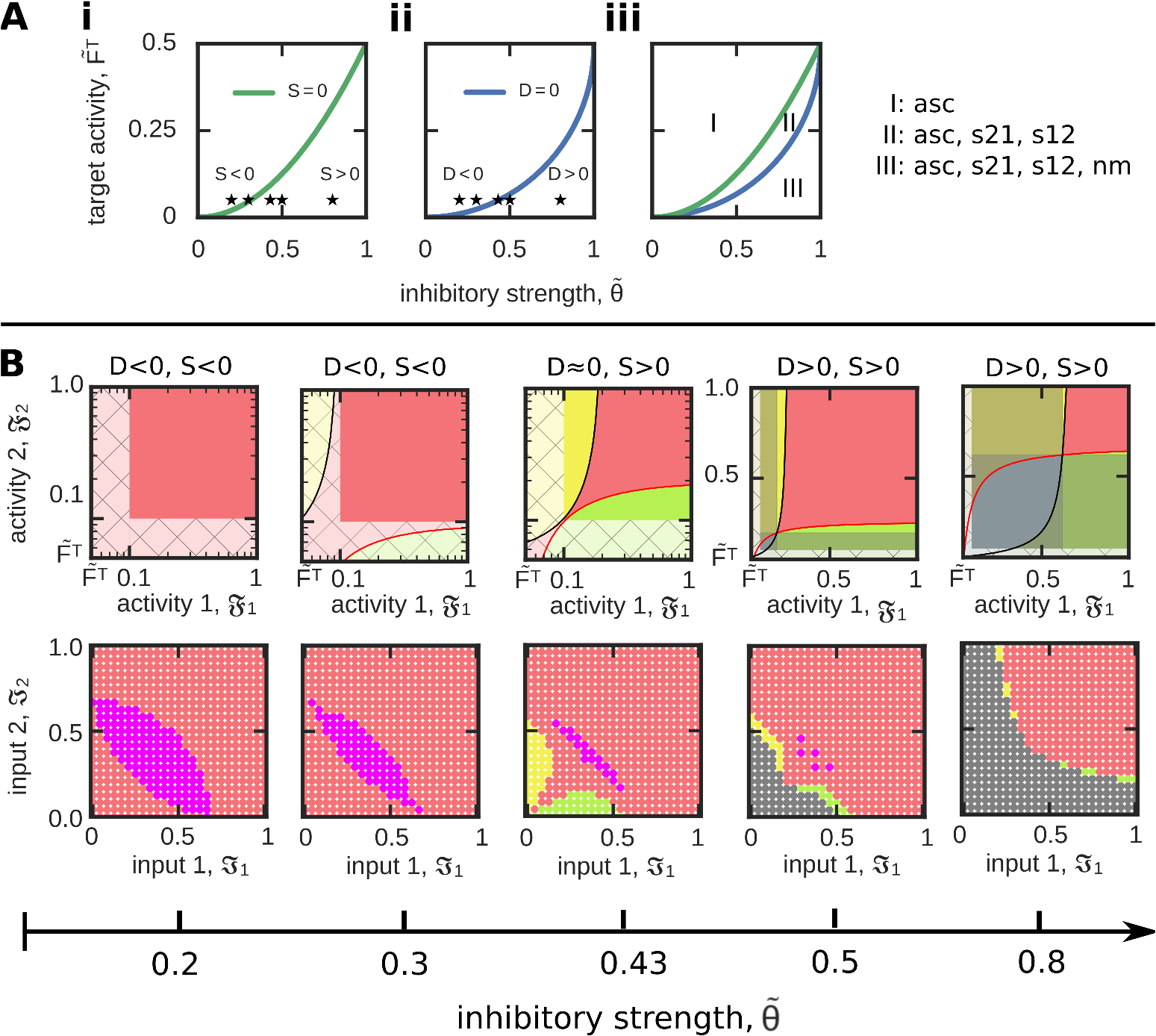
Quantification of the system ability to form different FOs dependent on the parameters 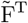 and 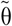. (**A**) The measure *S* (i; green) indicates whether sequences can be formed, while the measure *D* (ii; blue) specifies the existence of no-memory states. (iii) Both measures together separate the 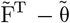–parameter phase space into three distinct regimes. Please see main text for details. (**B**) For a constant target activity 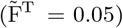, we show several examples of resulting functional organizations for different values of inhibition 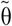 (asterisks in (A)) in activity- (top row) and input-space (bottom).

Thus, the 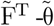-(dependency of *D* (Fig 4 A i) and *S* (Fig 4 A ii) determine the potential of the system to form diverse functional organizations (Fig 4 A iii). Interestingly, there are three functionally different system configurations: For *D* < 0, *S* < 0, the system can only form associations (regime I in Fig 4 A iii; first and second column in Fig 4 B). If *D* < 0, *S* > 0, the system can form either associations or sequences (*s12* as well as *s21*; regime II; third column). And if *D* > 0, *S* > 0, associations, sequences, and the no-memory state can be formed and reached by the system (regime III; fourth and fifth column). Thus, this analysis shows that with larger average inhibitory weight 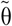 and smaller target activity level 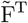 the system receives a larger repertoire of functional organizations. However, this analysis also shows that the functional organization of discrimination cannot be formed in a long-term manner. Although for large values of inhibition both separatrices intersect (see, for instance, fifth column in Fig 4 B), thus, both activity levels could be simultaneously below their corresponding separatrix (blue), the resulting area of discrimination cannot be reached by any inputs ℑ_1_, ℑ_2_ because in all these cases the neuronal populations cannot serve as memory representations (grey).

##### Theorem 2

Correlation-based synaptic plasticity in combination with a postsynaptic activity-dependent synaptic scaling term lacks the formation of functionally unrelated memories (discrimination).

##### Proof.

The constraints for a discrimination of two memories can be summarized by two inequations regarding the mean synaptic weights of the neuronal population model

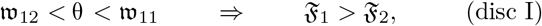

and

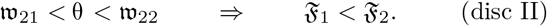

We easily see that condition disc I is in contradiction with condition disc II, and thus, the discrimination of two interconnected memories is excluded.

Thus, a neuronal system with correlation-based synaptic plasticity and a postsynaptic-activity-dependent synaptic scaling is not able to form two excitatory relations in between two memory representations which are weaker than the average inhibition. This analysis reveals that applying such a learning rule globally for the neuronal network dynamics is not sufficient to distinct the two different processes of memory formation and discrimination. Thus, it seems that this learning rule has to be augmented by at least one additional adaptive process that decouples these two processes.

### Local inhibition enables the functional organization of discrimination

The ability to form a discriminatory relation between memory representations is functionally very important for a neuronal system, as it implies that not all memories, which are anatomically connected with each other, have to be functionally connected with each other. Thus, to overcome the lack of discriminatory functional organizations of memories, we have to “decouple” the discrimination condition from the memory condition (see above).

For this, we introduce a different inhibitory synaptic weight strength 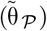 within the neuronal populations compared to the inhibitory synaptic weight strength for all other connections (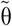, Fig 5 A). In other words, the parameter 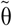 is different for the discrimination condition as for the memory condition (which is now 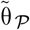). To quantify the influence of this new parameter on the potential to form two discriminated memory representations, we calculate the size of the activity space leading to discrimination (Fig 5 B, left). In general, if inhibition within the populations is weaker than for all other connections 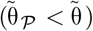, the system can form memories being in a discrimination (Fig 5 B, right). Please note that the other functional organizations are still maintained such that all different types can be obtained (Fig 5 C). While in this analysis we predefined different levels of inhibition, these different levels can also be obtained by the system in a self-organized manner by considering inhibitory synaptic plasticity (see Fig 6 for an example of discrimination). Similar to excitatory synaptic plasticity, the here-used inhibitory plasticity rule depends on the correlation of pre- and postsynaptic firing. In addition, the inhibitory synaptic plasticity rule is multiplied by two additional constrains. First, a minimum activity level θ_*F*_ for the pre- and postsynaptic firing rates introduces a threshold for inhibitory synaptic plasticity to occur

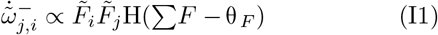

with 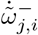 being the strength ofthe inhibitory synapse connecting the presynaptic neuron *i* with the postsynaptic neuron *j*, 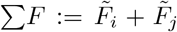, and H being the heaviside step function. Second, the difference in the pre- and postsynaptic firing rates 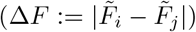 provides an abstract measure for the non-correlation of firing due to large deviations in their firing rates.

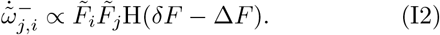

**Fig 5.**
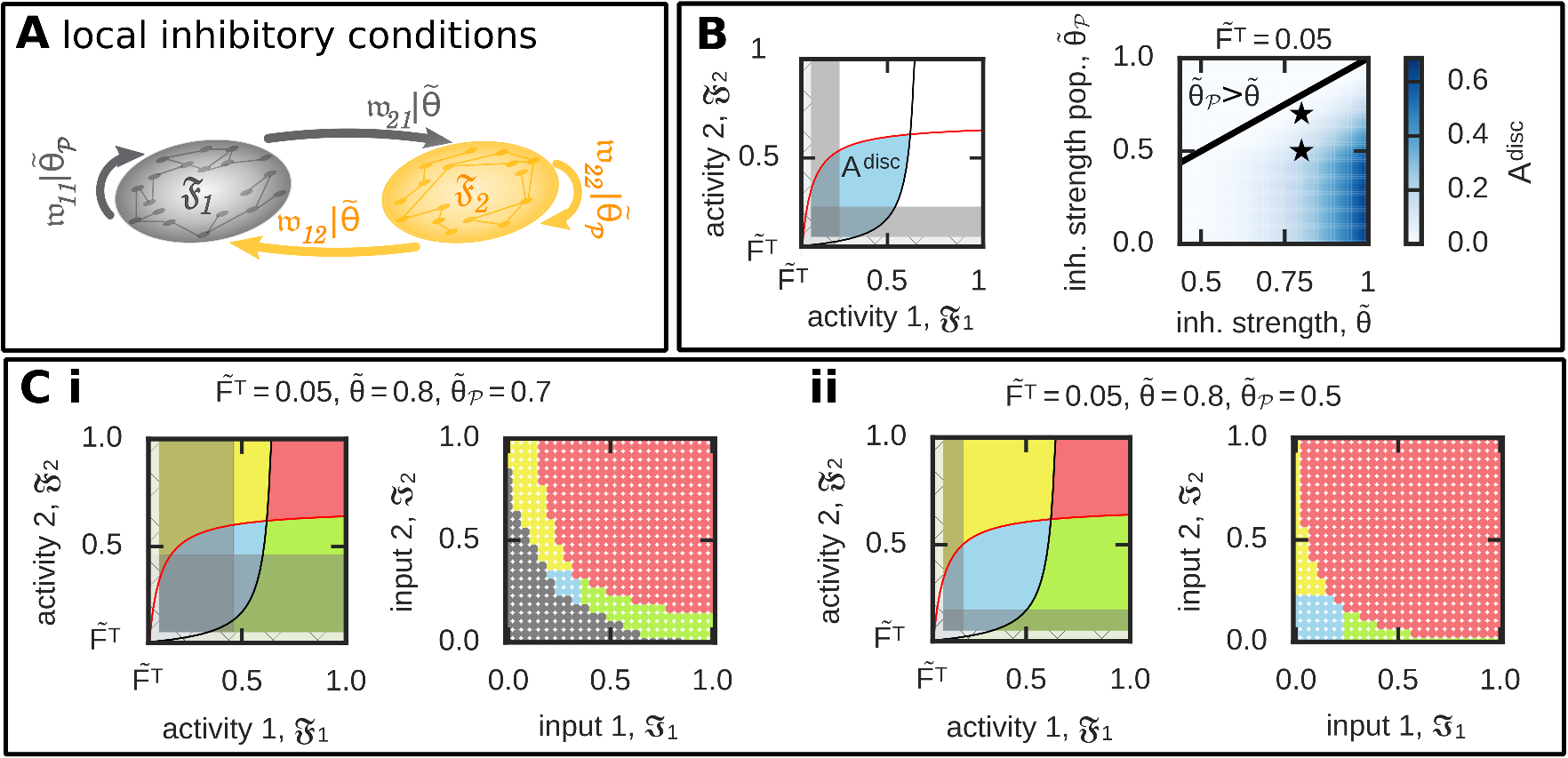
Considering different levels of inhibition level for connections within the neuronal populations compared to all others enables the formation of two discriminated memory representations. (**A**) We consider a different average inhibitory synaptic weight within the neuronal populations 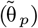 compared to all others 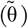. (**B**) Left: To quantify the effect of different inhibition levels, we calculate the area of discrimination states (A^disc^; blue) not being “covered” by the no-memory states (grey) in the 𝔉_1_-𝔉_2_-activity space. Right: A^disc^ dependency on different relations between 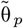 and 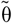. (**C**) Given a lower level of inhibition within the populations than otherwise provides the neural system the ability to form all functional organizations as indicated here by two examples (asterisked in (B)). 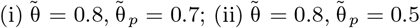.

**Fig 6.**
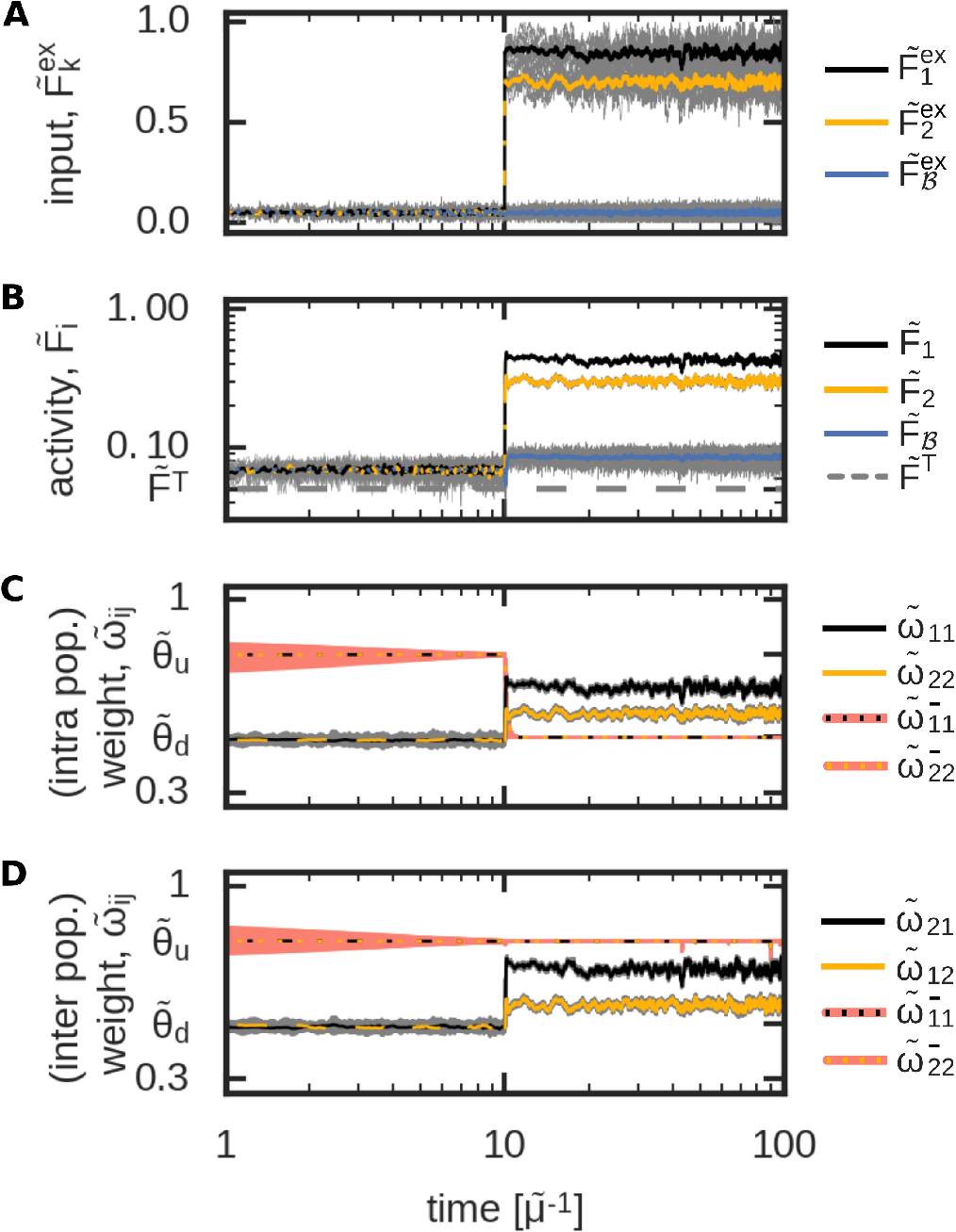
An exemplary inhibitory plasticity rule enables the self-organized formation of a discriminatory functional organization. The development of the input-driven dynamics of the complete neural network (Fig 1) with inhibitory plasticity. (**A**) The average input amplitudes are determined by two Ornstein-Uhlenbeck processes with mean 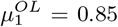 and 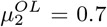. (**B**) Average activities of each population; (**C**) average intra-population excitatory and inhibitory synaptic weights; (**D**) average inter-population excitatory and inhibitory synaptic weights.

Here, *δF* describes a tolerance range for such a variation in the firing rates.

According to these conditions, the inhibitory synaptic weights converge either to

- an up-state (θ_*u*_), if the sum of neuronal activities is smaller than its threshold (Σ *F* < θ_F_) **and/or** the difference in the pre- and postsynaptic activities is above its tolerance range (Δ*F* > *δF*), or
- a down-state (θ_*d*_), if the sum of neuronal activities is larger than its threshold (Σ *F* > θ_*F*_ **and** the difference in the pre- and postsynaptic activities is smaller than its tolerance range (Δ*F* < *δF*).

This type of inhibitory synaptic plasticity together with plastic excitatory synapses governed by the interaction of correlation-based synaptic and homeostatic plasticity enables the reliable formation of memory representations and, in addition, provides the system the ability to form all basic functional organizations. In other words, our analysis indicate that a self-organized neural network can form all types of functional organizations if the interactionof synaptic plasticity and scaling are complemented by further adaptive processes.

## DISCUSSION

### General framework

In the present work, we have developed a mathematical framework to investigate the ability of adaptive neural networks to form in a dynamic, input-dependent manner diverse functional organizations of interconnected memories. In contrast to previous studies focusing only on a subset of possible functional organizations [15, 21, 25, 29–31], we consider here all possible organizations between two memory representations. Thereby, we define the functional organizations dependent on the relation between the excitatory and inhibitory synaptic weights of the neuronal network. By introducing a population description, we are able to transfer the resulting high-dimensional problem to a low dimensional problem considering average synaptic weights and activities of the neuronal populations involved. In addition, by considering the long-term equilibrium dynamics, we could further reduce the system complexity with the input stimulation being a system parameter. Finally, we could map the resulting dynamics onto the two-dimensional activity-space which is sufficient to solve this complex problem of memory interactions (Fig 2). Thus, we gain an easily accessible understanding of the possible states the system can reach as well as of the underlying principles arising from the considered plasticity mechanisms and their limitations. Given the generality of the complete framework, it can be commonly used to investigate the effect of diverse plasticity mechanisms on the formation and interaction between memory representations.

### Analysis of the interplay between synaptic plasticity and synaptic scaling

Given this general mathematical framework, we analyzed the effect of the interplay of correlation-based synaptic plasticity with homeostatic synaptic scaling on the formation of functional organizations of memory. This type of interplay is a quite general formulation of synaptic dynamics [18, 32], which is sufficient to form individual memory representations [11, 14]. We have shown that these types of mechanisms provide a neural network the ability to form several types of functional organization of memory representations such as sequences and associations (Figs 3 and 4). Furthermore, our method shows that correlation-based plasticity with scaling does not enable the formation of two stable memory representations being in a discriminated state.

This shortcoming is due to the purely correlation-based formulation of synaptic plasticity and, by this, mathematically couple the condition for memory formation with the condition of discrimination. Interestingly, the correlation-independent dynamics triggered by synaptic scaling are not sufficient to decouple the conditions. However, these dynamics enable the formation of sequences providing a further functional role of synaptic scaling besides synaptic stabilization [18, 19, 33] and homeostatic regulation of neuronal activities [28, 32].

Based on our results, we expect that similar mathematical models of synaptic dynamics, which consist of correlation-based plasticity and a homeostatic term dependent on the postsynaptic activity level (e.g., Oja’s rule [34] or BCM rule [35]), are also not able to form memory representations in a discriminated state. Thus, a further factor determining the synaptic dynamics of the network is required to enable the functional organization of discrimination.

### Local variations of inhibition

We have shown that local variations in the level of inhibition could serve as such factor enabling the discrimination between memory representations and other functional organizations (Fig 5). Thereby, the average inhibitory synaptic strength within a memory representation has to be weaker than all other inhibitory synaptic weights. This is in contrast to the idea of an inhibition, which balances the strong excitation within interconnected groups of neurons [12, 36]. However, despite the local differences in the balance of inhibition and excitation, the network-wide levels of excitation and inhibition can still be in a balanced state [37, 38]. Furthermore, this type of inhibitory weight structure could emerge from an Anti-Hebbian-like inhibitory plasticity rule as discovered in the memory-related Hippocampus [39].

### Possible extensions of synaptic dynamics

Besides inhibition, other mechanisms could be the additional factor yielding, together with correlation-based and homeostatic plasticity, all functional organizations. For instance, spike-timing-dependent triggered LTD ([40, 41]; in contrast to firing rate-dependent LTD [35, 42, 43]) could be a measure of uncorrelated spike trains decoupling the memory from the discrimination condition. In more detail, the LTP-part of STDP [40, 44, 45] can be interpreted as a measure of the probability that the pre- and postsynaptic neurons fire correlated spikes during a small time window [17] described here in the rate-model by the correlation-based LTP-term. Whereas, the amount of uncorrelated spike pairs triggering LTD could be described in the here-used rate-model by the difference between the pre- and postsynaptic firing rates. We expect that considering such a difference-term would be sufficient to enable the formation of memory representations (by correlation-based LTP) in a discrimination state (by non-correlation-based LTD). This has to be verified in subsequent studies.

Please note that there is a multitude of studies indicating the existance of additional factors influencing synaptic plasticity. For instance, neuromodulatory transmitters, such as acetylcholine, noradrenaline, serotonin, and dopamine, can serves as third factor [46, 47]. However, with the mathematical framework developed in this study, it is now possible to investigate in more detail the effect of such factors on the formation, maintenance, and organization of memory representations in neuronal circuits. Furthermore, given the understanding of the organization between two memory representations, now one can extend this framework to investigate the self-organized formation of webs of memories and the emergence of complex behavior.

## MATERIALS AND METHODS

### Neuronal Network Model

We consider a recurrent neuronal network model consisting of a set 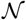 of *n* rate coded neurons 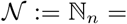 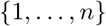, Fig 1 A, dots). The neurons are interconnected via an all-to-all connectivity for the excitatory as well as for the inhibitory connections. Note that, if not stated otherwise, the inhibitory connections are constant while the excitatory synapses are plastic. Within the recurrent network, we define two distinct subsets of neurons 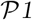 and 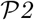 as *neural population 1* (black dots) and *neural population 2* (yellow dots). For simplicity, both neural populations have the same number of neurons 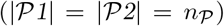. Furthermore, we assume no overlap between both neuronal populations 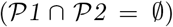. Neurons which are not part of neuronal population 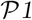 or 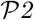 are summarized as *background neurons* 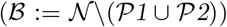, with size 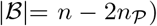. Thus, we can describe the neuronal network model as the interaction of three different neuronal populations 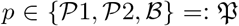. All neurons *i* of a neuronal population *p* receive a population-specific input stimulation 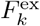 via 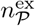 different neurons *k* from the input layer *ε*_*i*_ via constant excitatory synapses ω^ex^. Thus each neuron *k* of the external input layer provides an external input stimulus of average strength 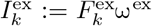 onto the interconnected neurons of the neuronal network 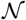. Note, we set 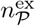 equal to 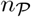 to consider the same order of magnitude for input populations as for the populations themselves.

### Neuron Model

We consider point-neurons with each neuron 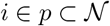 of the network summing up its incoming inputs from the interconnected excitatory (Fig 1 A, blue connections), inhibitory, and input neurons (red connections) to its overall neuron-specific input current (*ϕ*_*i*_). The inputs are transmitted via the synapses, thus, the neuron specific input current integrates the separate inputs of the neurons (*F*_*j*_) proportional to the respective synaptic weights (ω_*i*,*j*_):

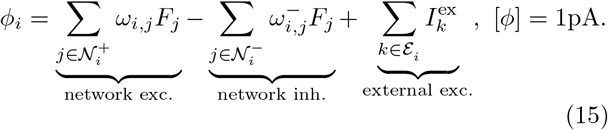

Here, 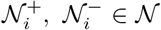 are the sets of indices for the excitatory, inhibitory, and externally interconnected presynaptic neurons *j* to the postsynaptic neuron *i*, whereby, ω_*i*,*j*_ and 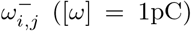 are the respective synaptic weights, and *F*_*j*_ ([*F*] = 1s^−1^) the respective presynaptic neuron’s activity. Due to the all-to-all connectivity of the network, the sets for the excitatory and inhibitory interconnected neurons equal the set of the neuronal network 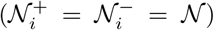. In contrast to the plastic excitatory synaptic weights, all inhibitory synapses 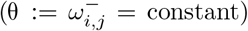 are constant. This leads to the following input current (*ϕ*_*i*_) onto one neuron *i* belonging to population *p*:

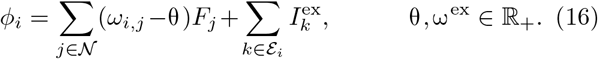

We can further specify this input current in respect to the presynaptic neuron’s affiliation to a neuronal population 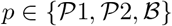:

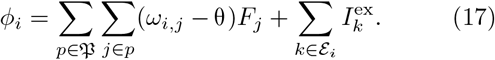

This neuron specific input, drives its respective membrane potential (*u*_*i*_) described by:

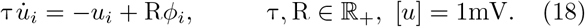

Here, τ ([τ] = 1s) is the time constant for the membrane potential and set to τ = 1s and R = 0.1nΩ, ([R] = 1nΩ) is the membrane resistance. The membrane potential *u*_*i*_ is non-linearly transformed to a neural firing rate (*F*_*i*_):

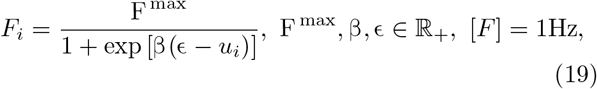

with F^max^ = 100Hz being the maximal firing rate, β ([β] = 1mV^−1^) being the steepness and ϵ ([ϵ] = 1mV) being the inflexion point of the sigmoid. Thus, the neuronal activity for each neuron *i* takes values between 0 and F ^max^ (*F*_*i*_ ∈ [0, F ^max^]). To simplify the description of the neuronal dynamics, we combine Eq 18 and Eq 19 to:

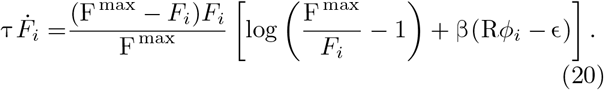

### Synaptic Plasticity and Synaptic Scaling

All excitatory synapses within the recurrent network are plastic and change proportional to the activity-dependent Hebbian learning rule (*H*, [3, 48])

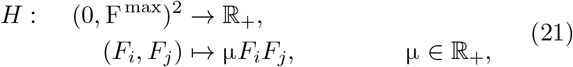

with time constant μ. This correlation learning rule lead to unbounded synaptic weight dynamics. Thus, we include synaptic scaling (*S*, [16, 28]) as a homeostatic mechanism

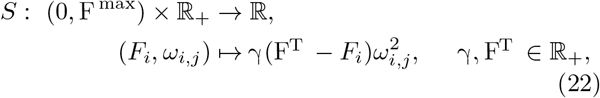

with the time constant γ and target firing rate F^T^. This scaling mechanism decreases (increases) the synaptic weight if the postsynaptic activity is above (below) the target firing rate. Note that Hebbian dynamics are in general faster than scaling dynamics, thus, γ ≪ *μ* Therefore, we set 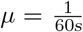 on the time scale of minutes while 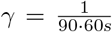 on time scale of hours. Combining the correlation based Hebbian learning term additive with the postsynaptic activity-dependent synaptic scaling term 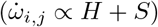, we get the following learning rule for the synaptic weights [18, 27]:

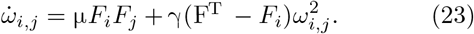

### Constraining parameters of the activity function

In the following, we give an interpretation for the parameters like the inflexion point and steepness of the sigmoidal shaped activity function (Eq 19).

To specify the inflexion point of the neuronal activities, we first, define the maximal evoked membrane potential (*u*^max^) of a neuron *i* by only one incoming synapse (ω_*ij*_) to the postsynaptic neuron *j*. Therefore, we set the pre- and postsynaptic neuronal activities to the maximal activity level of (*F*_*j*_ = *F*_*i*_ = *F*^max^) and by this calculate the fixed synaptic weight, using Eq (23), and define it as the maximal synaptic weight 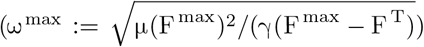. Eq 18, by this, specifies the maximal network internal 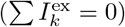 evoked membrane potential of *u*^max^:= RF^max^(ω^max^ − θ). Using this quality of *u*^max^, we interpret the inflexion ϵ of a neuron *i* as the number (*n*_ϵ_) of such maximally wired presynaptic neurons. This leads to ϵ = *n*_ϵ_*u*^max^. For the determination of the precise value for *n*_ϵ_ = 12 see the Results section.

To specify the steepness of the neuronal activity function, we have to consider their maximal and minimal possible evocable membrane potential and choose a steepness parameter β due to two different constraints: (i) the activity for the minimal membrane potential has to take on higher values as the target firing rate F^T^ to prevent unstable weight dynamics, and, (ii) for maximal evoked membrane potential the neurons have to take on the maximal firing rate of F ^max^. One specific parameter for the steepness of the activity function that full fill these two conditions is β = 0.00035mV^−1^ for all neurons.

### Normalized Neuronal Network Model

In the following, to reduce complexity, we normalize the neuronal activities of all neurons 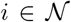 according to the maximal neural firing rate 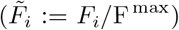 and all synaptic weights to the maximal excitatory synaptic weight 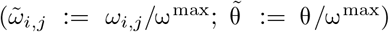. Thus, the external input stimulation is also normalized to 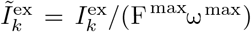. By this, we map the (normalized) neuronal activity 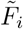 and the (normalized) excitatory synaptic weight 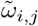 to [0,1] ∈ ℝ:

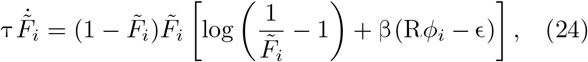

for the neuronal activity with

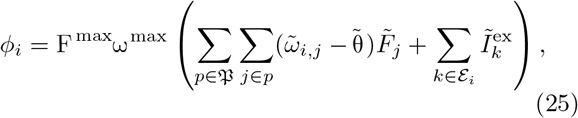

and

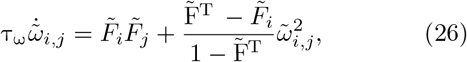

for the synaptic weight with

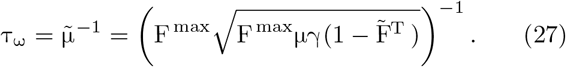

### Numerical Simulation and Stimulation Protocol

Each neuronal simulation starts with a tuning phase, where all neurons of the network receive input noise from 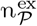 different input neurons of the input layer ε. Subsequent to this tuning phase (*t* = 10), all neurons *k* of the external input layer ε_*i*_ that are connected to neurons *i* ∈ *p* fire according to an Ornstein-Uhlenbeck process:

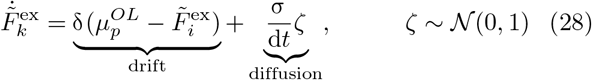

with an drift term with constant δ = 0.025 and an arbitrary population specific equilibrium level for the firing rate of 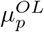, a normal distributed diffusion term with constant σ = 0.0125 and an initial firing rate of 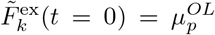. We apply this input stimulation over time, by this, simulate the whole activity and synaptic weight dynamics until the system reaches an equilibrium state. Numerically, we solve the differential equations of the normalized model for the synaptic weight dynamics and activity dynamics with the *euler method*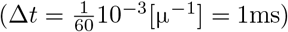.

### Analysis of the System’s Equilibrium State

As the different functional organizations of two interconnected neuronal populations are defined at the system’s equilibrium state, we can reduce our problem to an analytically calculation of the average neuronal activities and synaptic weights at equilibrium state. Due to the homogeneous external input stimulation of all neurons *i* belonging to one neuronal population *p* and the underlying full connectivity of the network, we make the assumption that the fixed firing rate of each neuron *i* of a population 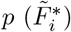 approaches the mean firing rate of the particular population (𝔉_*p*_) at fixed point state

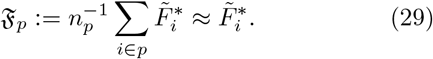

Using these average activities (𝔉_*p*_) of both neuronal populations *p* at stable state we can calculate the respective average excitatory synaptic weights at the system’s stable state

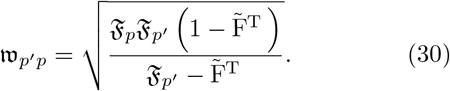

This approach, reduces the problem to analytically calculate the average activities (𝔉_*p*_) of the neuronal populations 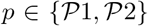 dependent on the external input stimulation. Therefore, we calculate the mean population specific external input current onto one neuron *i* of population p as 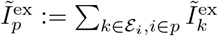.

We easily see that the input current onto each neuron *i* of a neuronal population *p* is independent on the neuronal dynamics and is only defined by the average qualities of each neuronal population at the system’s stable state. Thus the fixed mean input onto a neuronal population *p* (𝔔_*p*_) is given by:

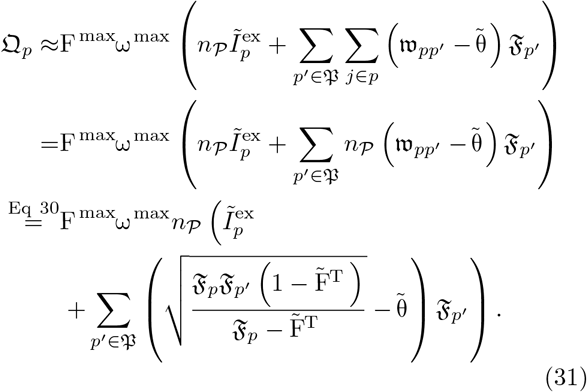

We further reduce the complexity of the model to enable a fixed point analysis, while adding the fixed input from the background neurons onto one neuronal population *p* at stable state

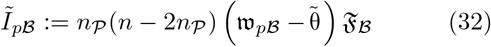

to the external input stimulation 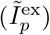, proportionally to the size of the external input layer 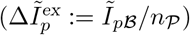. This leads to the total input current onto one neuronal population 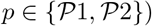, as:

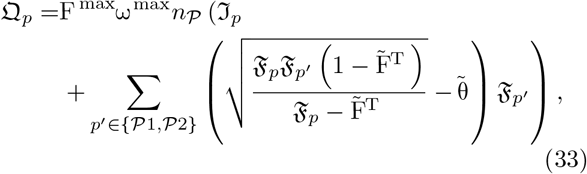

with 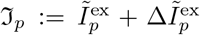. This expression of the input current onto each neuronal population *p* at stable state, allows us to numerically calculate the respective average activity of each population at the system’s stable state:

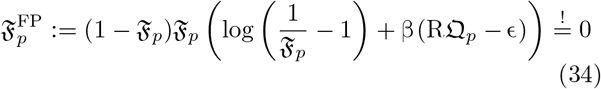

in a two dimensional parameter-phase space of 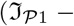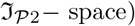.

### Inhibitory synaptic plasticity

In the last section of this present work, we introduce inhibitory synaptic plasticity. This specific plasticity rule depend on a threshold (θ_*F*_) for the sum of pre- and postsynaptic activity levels 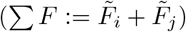 and a tolerance range (*δF*) for the difference in the pre- and postsynaptic firing rates 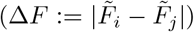 leading the inhibitory synaptic weight 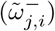 converge either to an up- (θ_*u*_) or down-state (θ**d**). The synaptic weight converge to:

- an up-state (θ_*u*_), if the sum of neuronal activities is smaller than its threshold (Σ *F* < θ _*F*_) **and/or** the difference in the pre- and postsynaptic activities increases its tolerance range (Δ*F* > *δF*)
- a down-state (θ_*d*_), if the sum of neuronal activities is greater than its threshold (Σ *F* > θ _*F*_) **and** the difference in the pre- and postsynaptic activities is smaller than its tolerance range (Δ*F* < *δF*).

These conditions lead to the following learning rule on the inhibitory synaptic plasticity

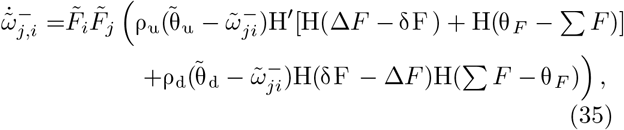

with ρ_*u*_, ρ_*d*_ being the learning rates towards the up- and down-state and H′ an adapted heaviside function with H′(0) = 0 to express the and/or condition for the up-state. In our simulations we set θ_*u*_ = 0.8, θ_*d*_ = 0.5, θ_F_ = 2𝔉^min^, *δF* = 0.05 and ρ_*d*_ = ρ_*u*_ = 1.

## ACKNOWLEDGMENTS

We thank Florentin Wörgötter for fruitful comments. The research was funded by the H2020-FETPROACT project Plan4Act (#732266) [JH, CT] and by the International Max Planck Research School for Physics of Biological and Complex Systems by stipends of the country of Lower Saxony with funds from the initiative “Niedersächsisches Vorab” and of the University of Göttingen [JH].

